# Chemogenetic stimulation of mouse central amygdala corticotropin-releasing factor neurons: Effects on cellular and behavioral correlates of alcohol dependence

**DOI:** 10.1101/2020.02.07.939496

**Authors:** Max Kreifeldt, Melissa A Herman, Harpreet Sidhu, Giovana C de Macedo, Roxana Shahryari, Marisa Roberto, Candice Contet

## Abstract

**Background:** Corticotropin-releasing factor (CRF) signaling in the central nucleus of the amygdala (CeA) plays a critical role in rodent models of excessive alcohol drinking. However, the source of CRF acting in the CeA during alcohol withdrawal remains to be identified. In the present study, we hypothesized that CeA CRF interneurons may represent a behaviorally relevant source of CRF to the CeA increasing motivation for alcohol via negative reinforcement.

**Methods:** We tested this hypothesis in male mice and used chemogenetics to stimulate CeA CRF neurons in vitro and in vivo.

**Results:** We first observed that Crh mRNA expression in the anterior part of the mouse CeA, at the junction with the interstitial nucleus of the posterior limb of the anterior commissure, correlates positively with alcohol intake in C57BL/6J males with a history of chronic binge drinking. We then found that chemogenetic activation of CeA CRF neurons in Crh-IRES-Cre mouse brain slices increases gamma-aminobutyric acid (GABA) release in the medial CeA in part via CRF1 receptor activation, indicating local CRF release. While chemogenetic stimulation of CeA CRF neurons exacerbated novelty-induced feeding suppression, as seen in C57BL/6J males withdrawn from chronic intermittent alcohol inhalation, it had no effect on voluntary alcohol consumption, following either acute or chronic manipulation.

**Conclusions:** Altogether, these findings indicate that hyperactivity of CeA CRF neurons may contribute to elevated CeA GABA levels and negative affect during alcohol withdrawal but is not sufficient to drive alcohol intake escalation in dependent mice.

## Introduction

Alcohol use disorders (AUDs) represent a spectrum of pathological patterns of alcoholic beverage consumption that affect 6.2% of the adult population in the United States of America [1] and more than 100 million people worldwide [2]. AUDs are not only characterized by the intake of excessive amounts of alcohol but also by the emergence of a negative emotional state (e.g., anxiety, irritability) upon withdrawal. While the motivation to drink alcohol is initially driven by the desire to experience the pleasurable effects of intoxication (positive reinforcement), it becomes progressively driven by the need for relief from the negative emotional state associated with withdrawal (negative reinforcement) [3]. Excessive alcohol drinking (i.e., beyond intoxication threshold) and negative affect during withdrawal can be triggered in several mouse and rat models, which has enabled the dissection of underlying neurobiological mechanisms [4–8].

Multiple pharmacological and genetic studies have highlighted that recruitment of the extrahypothalamic corticotropin-releasing factor (CRF, encoded by the *Crh* gene) system plays a pivotal role in the emergence of negative reinforcement in alcohol-withdrawn animals, whereby activation of CRF type 1 receptors (CRF1) contributes to the excessive drinking, heightened anxiety and stress sensitization characterizing alcohol dependence (see [3, 9–11] for review). In particular, CRF is released in the central nucleus of the amygdala (CeA) during alcohol withdrawal [12] and blockade of CRF1 signaling in the CeA reduces the anxiogenic-like effect of alcohol withdrawal [13, 14], dependence-induced increases in alcohol self-administration [15, 16], as well as heavy binge drinking [17]. CRF1 in the CeA also mediates the elevation of gamma-aminobutyric acid (GABA) dialysate induced by alcohol dependence [18] and CRF1 overexpression in the CeA potentiates stress-induced reinstatement of alcohol seeking [19].

Despite this wealth of converging evidence, the specific neurons responsible for the release of CRF in the CeA during alcohol withdrawal have not been identified. In the present study, we tested the hypothesis that these neurons are intrinsic to the CeA. This hypothesis was supported by evidence that *Crh* is upregulated in the CeA of alcohol-dependent and post-dependent rats [18, 20, 21], along with our own findings (reported here) that *Crh* expression in the anterior part of the CeA correlates with alcohol intake in mice with a history of chronic alcohol drinking followed by abstinence. Importantly, psychological stress also upregulates *Crh* in the CeA [22–24] and CRF synthesis in the CeA plays a critical role in mediating anxiety-like behavior [25, 26]. In addition, CRF1-expressing neurons in the CeA receive direct input from local interneurons [27] and CeA CRF neurons provide both local inhibitory GABA and excitatory CRF1-mediated signals to other CeA neurons in *Crh*-Cre rats [28], which supports the notion that CRF can be released from intrinsic CeA neurons. Furthermore, studies in *Crh*-Cre rats have recently shown that chemogenetic stimulation of CeA CRF neurons elicits anxiety-like behavior [29, 30], while their optogenetic inhibition reduces escalated alcohol self-administration in alcohol-dependent rats [31].

According to the hypothesis that CeA CRF neurons can release CRF locally, we predicted that chemogenetic stimulation of mouse CeA CRF neurons would replicate cellular and behavioral hallmarks of alcohol dependence. While this manipulation produced local CRF1-mediated GABA release and elicited signs of negative affect resembling those seen during alcohol withdrawal, it did not increase voluntary alcohol consumption.

## Materials and Methods

### Animals

C57BL/6J males were purchased from the Jackson Laboratories (stock #000664). *Crh*-IRES-Cre male breeders were obtained from The Jackson Laboratory (B6(Cg)Crh^tm1(cre)Zjh^/J, stock # 012704, [32]) and were mated with C57BL/6J females from The Scripps Research Institute rodent breeding colony to generate the heterozygous males used for experimentation. Additional details are provided in the Supplement. All procedures adhered to the National Institutes of Health Guide for the Care and Use of Laboratory Animals and were approved by the Institutional Animal Care and Use Committee of The Scripps Research Institute.

### Viral vectors

Adeno-associated viral serotype 2 (AAV2) vectors encoding the hM3Dq excitatory designer receptor [33] fused to the red fluorescent protein mCherry, under the control of the human synapsin promoter and in a Cre-dependent manner (AAV2-hSyn-DIO-hM3Dq-mCherry [34]), were obtained from the Vector Core at the University of North Carolina (UNC) at Chapel Hill or from Addgene. Vectors were injected bilaterally into the anterior part of the CeA (AP −0.9 mm from bregma, ML ± 3.0 from the midline, DV −4.5 mm from the skull). Additional details are provided in the Supplement.

### Experimental cohorts

*In situ* hybridization data were collected from a cohort of C57BL/6J mice subjected to chronic alcohol drinking (n=15). For chemogenetic experiments, a first cohort of *Crh*-IRES-Cre mice was used for electrophysiological recordings (n=12). A second cohort was tested in the elevated plus-maze (EPM), social approach, and novelty-suppressed feeding assays, with at least one week between tests; these mice were then single-housed and tested for alcohol drinking (n=14). A third cohort was tested for digging and marble burying, and their brains were used to quantify c-Fos induction one week later (n=15). A fourth cohort was used to confirm the phenotype observed in the novelty-suppressed feeding test (n=12) and include negative control mice (n=8); these mice were also tested for locomotor activity and fasting-refeeding in the home cage.

Detailed methods for *in situ* hybridization, electrophysiology, immunohistochemistry and behavioral testing are provided in the Supplement.

### Statistical analysis

Data analysis was performed in Statistica 13.3 (TIBCO Software Inc.). The correlation of *Crh in situ* hybridization signal with ethanol intake was evaluated using Pearson’s r. The effect of clozapine-N-oxide (CNO) on firing rate and sIPSC parameters was analyzed using paired two-tailed Student’s t-tests, and the effect of CNO combined with R121919 was analyzed by repeated-measures analysis of variance (RM-ANOVA). RM-ANOVAs were also used to analyze the effect of CNO on ethanol intake (acute regimen) and refeeding. The effects of CNO on ethanol intake (chronic regimen), social approach and hyponeophagia were analyzed by two-way RM-ANOVA. Bonferroni *posthoc* tests were conducted when relevant. The effects of CNO on c-Fos counts, hyponeophagia replication (pre-planned comparisons) and other behavioral endpoints were analyzed using unpaired two-tailed Student’s t-tests. Data are shown as mean ± s.e.m. on the graphs.

## Results

### Crh expression in the anterior part of the CeA correlates with ethanol intake in chronic binge drinking mice

As described previously using CRF immunostaining and a GFP reporter strain, CRF expression varies along the rostro-caudal extent of the mouse CeA [35, 36]. We confirmed this observation using chromogenic *in situ* hybridization to label *Crh* mRNA (Fig. 1A), as can also be seen in the Allen Mouse Brain Atlas (https://mouse.brain-map.org, experiment 292). At the most anterior level (≈ −0.8 mm from bregma), a V-shaped cluster of CRF neurons is found at the transition between the interstitial nucleus of the posterior limb of the anterior commissure (IPAC) and the CeA. At midlevel (≈ −1.2 mm from bregma), CRF neurons are more sparsely distributed, mostly within the lateral aspect of the CeA, and to a smaller extent within the medial CeA. At a more posterior level (≈ −1.6 mm from bregma), CRF neurons are tightly packed within the lateral CeA, with again a few cell bodies found in the medial CeA.

**Figure 1.**
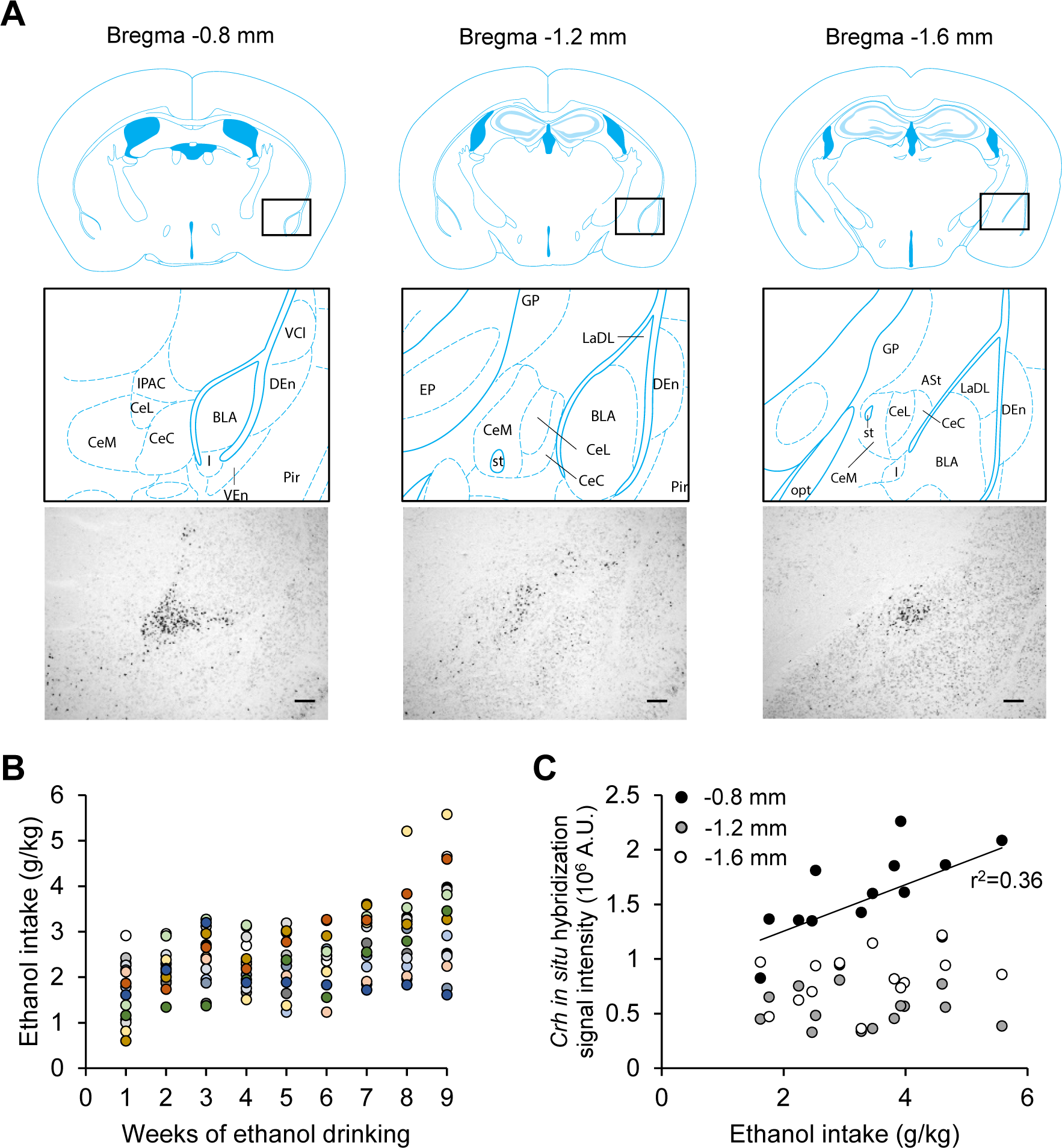
*Crh* mRNA levels in the anterior CeA correlate with ethanol intake in mice subjected to chronic binge drinking. **A.** Representative images of *Crh* mRNA distribution at three antero-posterior levels of the mouse CeA (scale bars = 200 μm) and corresponding brain atlas diagrams highlighting a V-shaped cluster of CRF neurons at the junction of the anterior CeL and IPAC (left panels), and scattered CRF neurons at more posterior levels of the CeL (middle and right panels). Brain atlas diagrams were reproduced from [67]. BLA, basolateral amygdaloid nucleus; CeC, capsular part of the CeA; CeL, lateral division of the CeA; CeM, medial division of the CeA; DEn, dorsal endopiriform claustrum; EP, entopeduncular nucleus; IPAC, interstitial nucleus of the posterior limb of the anterior commissure; GP, globus pallidus; I, intercalated nuclei of the amygdala; LaDL, lateral amygdaloid nucleus, dorsolateral part; opt, optic tract; Pir, piriform cortex; st, stria terminalis; VCl, ventral part of claustrum; VEn, ventral endopiriform claustrum. **B.** Average weekly ethanol intake in a cohort of mice subjected to 2-h two-bottle choice (ethanol 15% v:v vs water) sessions five days per week for nine weeks. Each color represents an individual mouse. **C.** *Crh* chromogenic *in situ* hybridization signal density at three antero-posterior levels of the CeA as a function of ethanol intake during the last week. There was a significant correlation at the most anterior level (≈ bregma −0.8 mm), but not at more posterior levels.

Semi-quantitative measures of *Crh* expression at these three antero-posterior levels were obtained in C57Bl/6J males subjected to chronic alcohol binge drinking for nine weeks and euthanized four weeks after their last drinking session. Average ethanol intake during the last drinking week ranged between 1.6 and 5.8 g/kg per 2-h session (Fig. 1B). *Crh* expression in the most anterior part of the CeA (bregma −0.8 mm, including IPAC) correlated positively with the average ethanol intake measured during the last week of drinking (r^2^=0.36, p=0.02, Fig. 1C). No significant correlation was observed at more posterior levels of the CeA (bregma −1.2 mm: r^2^=0.02, p=0.65; bregma −1.6 mm: r^2^=0.11, p=0.23; Fig. 1C).

Based on these results, the anterior level of the CeA was targeted for viral vector infusions in subsequent chemogenetic experiments.

### Validation of chemogenetic approach to stimulate CeA CRF neurons

We used Cre-dependent expression of the excitatory designer receptor hM3Dq in *Crh*-IRES-Cre males to analyze the effects of CeA CRF neuron stimulation on CRF release, alcohol drinking and affective behaviors. We first assessed the specificity of Cre activity distribution in the CeA of Crh-IRES-Cre mice and verified that CNO activated hM3Dq-expressing cells both *in vivo* and in brain slices.

We analyzed the overlap between *Crh* and *tdTomato* mRNAs in CeA sections of Crh-IRES-Cre;Ai9 mice by double *in situ* hybridization (Fig. 2A). We found that 86.3 ± 4.9 % of *tdTomato*+ neurons expressed *Crh* (a measure of reporter fidelity), while 64.0 ± 5.4 % of *Crh*+ neurons expressed *tdTomato* (a measure of reporter penetrance).

**Figure 2.**
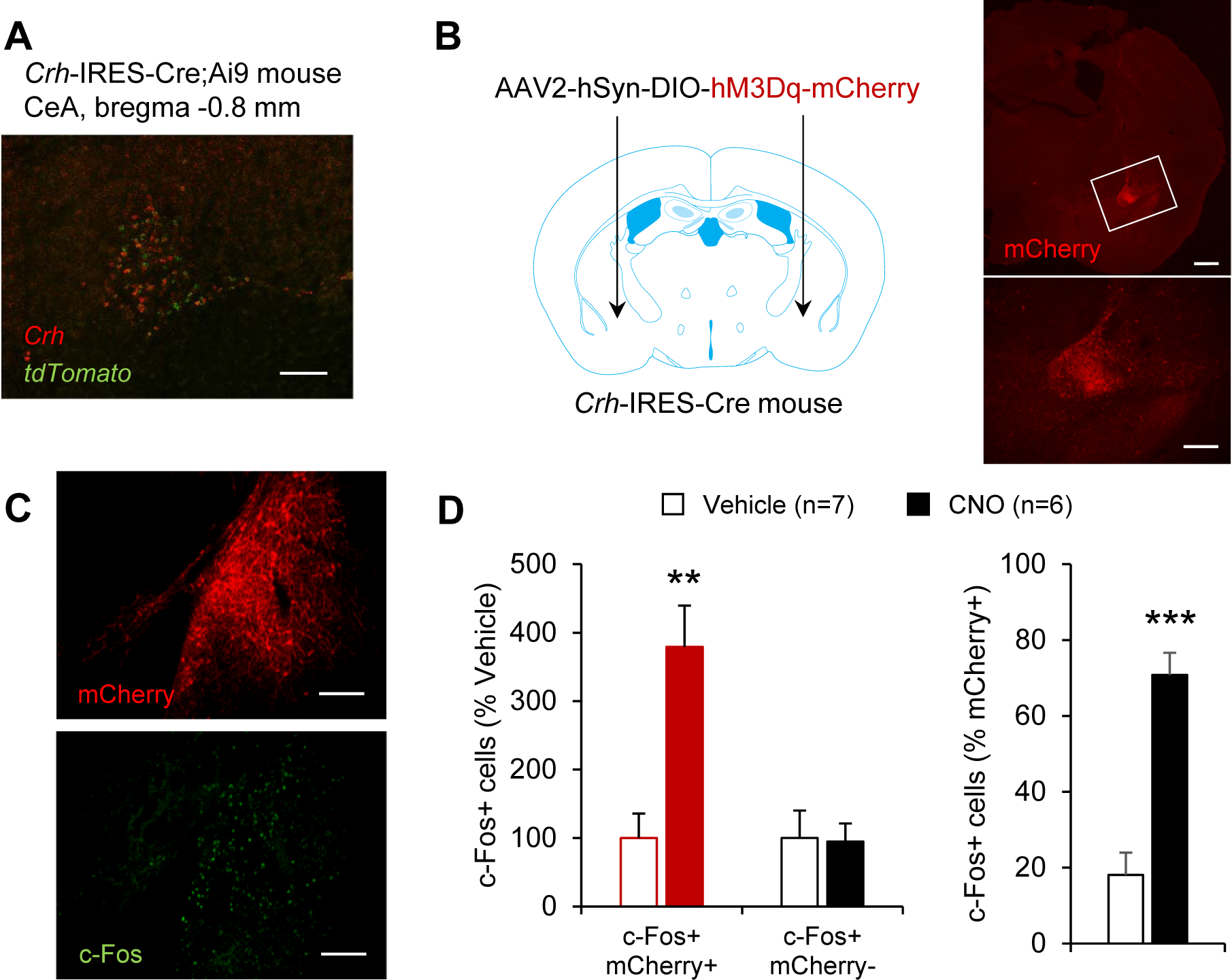
Validation of Cre activity and chemogenetic stimulation in CeA CRF neurons of *Crh*-IRES-Cre mice. **A.** Distribution of *Crh* (red) and tdTomato (green) mRNAs in the CeA of *Crh*-IRES-Cre;Ai9 mice, as visualized by double fluorescent *in situ* hybridization (scale bar = 100 μm). **B.** mCherry immunolabeling recapitulates the V-shaped pattern of *Crh* expression in the CeA of *Crh*-IRES-Cre mice injected with a Cre-dependent AAV vector encoding hM3Dq-mCherry. The area framed in the top picture (scale bar = 500 μm) in shown at higher magnification in the bottom picture (scale bar = 100 μm). **C.** Double immunostaining of mCherry (red) and c-Fos (green) was used to evaluate neuronal activation in the CeA following i.p. injection of vehicle (n=7) or CNO 5 mg/kg (n=6) in *Crh*-IRES-Cre mice injected with AAV2-hSyn-DIO-hM3Dq-mCherry in the CeA (scale bars = 100 μm). **D.** Chemogenetic stimulation of CeA CRF neurons increased c-Fos expression selectively in mCherry-positive CeA neurons. Data are shown as mean ± s.e.m. of the number of c-Fos+ cells expressed as percentage of Vehicle values (left graph) or percentage of mCherry+ cells (right graph). Data were analyzed by unpaired t-test: **, p<0.01; ***, p<0.001.

Consistent with these results, in *Crh*-IRES-Cre mice injected with an AAV2-hSyn-DIO-hM3Dq-mCherry vector in the CeA, mCherry-immunolabeled cell bodies showed the same distribution as *Crh* mRNA (Fig. 2B, compare with Fig. 1A left panel and Fig. 2A). Mice were perfused 90 min following i.p. administration of either vehicle or CNO (5 mg/kg) and their brains were processed for immunohistochemistry. mCherry immunoreactivity was used to label hM3Dq-expressing cells and c-Fos immunoreactivity was used as a marker of cellular activation (Fig. 2C-D). As expected, CNO increased the number of c-Fos+ cells selectively in the subpopulation expressing hM3Dq (t_11_=4.1, p=0.002), but it did not affect the number of c-Fos+ cells among mCherry-negative cells (t_11_=-0.1, p=0.91; Fig. 2D, left). Upon CNO administration, the proportion of active (c-Fos+) hM3Dq-expressing cells increased about 4-fold (t_11_=6.3, p<0.0001; Fig. 2D, right). In another cohort of *Crh*-IRES-Cre mice injected in the CeA with a Cre-dependent vector encoding hM3Dq-mCherry, current-clamp recordings of action potentials were obtained from mCherry-positive (i.e., hM3Dq-expressing) CeA neurons (Fig. 3A-B). As expected, CNO (500 nM) application significantly increased their firing rate (t_7_=-6.5, p=0.0003; Fig. 3B).

**Figure 3.**
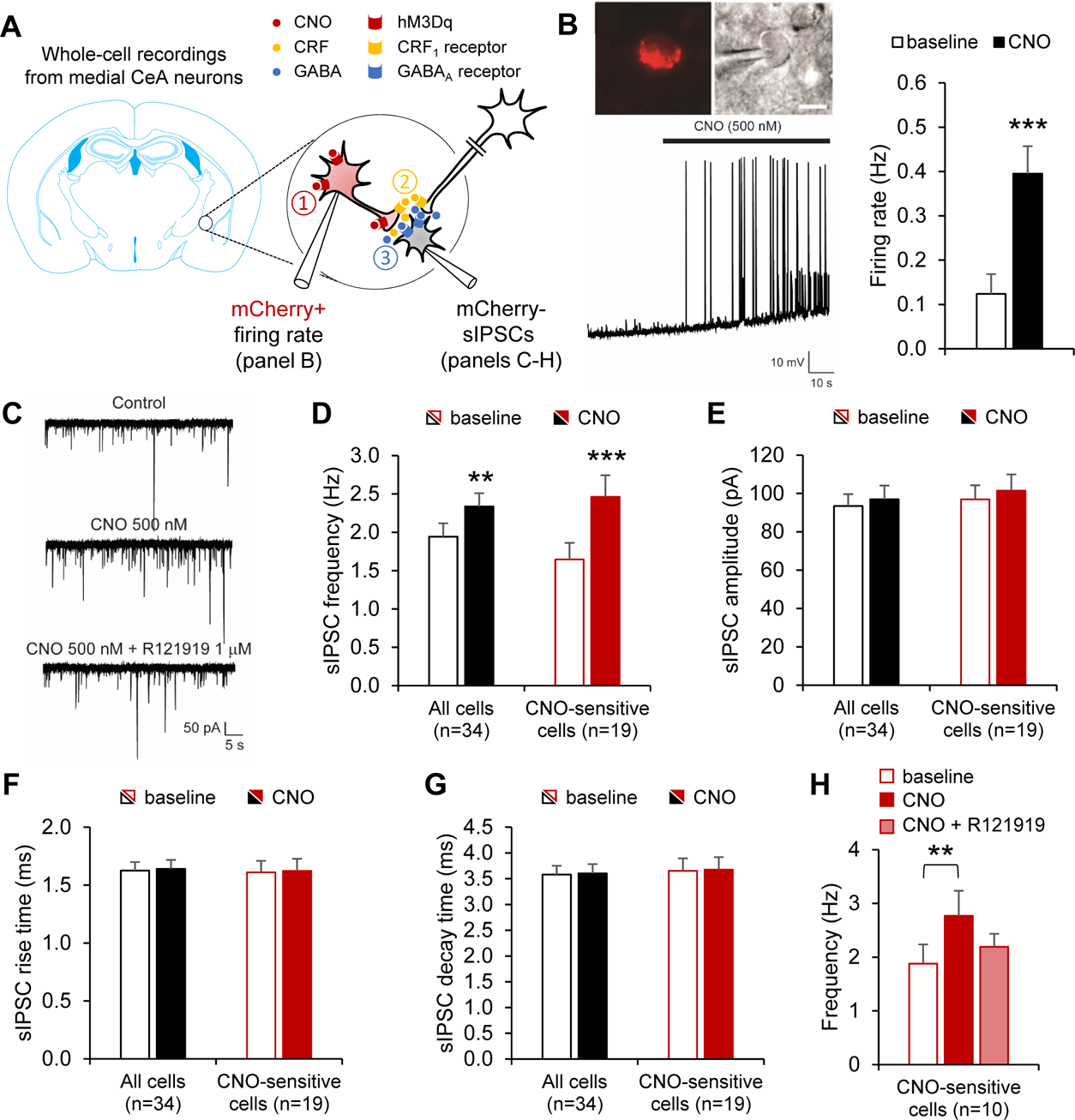
Chemogenetic stimulation of CeA CRF increases GABA release onto medial CeA neurons in a CRF1-dependent manner. **A.** *Crh*-IRES-Cre mice were injected in the anterior CeA with a Cre-dependent AAV vector encoding hM3Dq-mCherry. Whole-cell recordings were then obtained from mCherry+ neurons (panel B) or mCherry-neurons (panels C-H) to test the hypothesis that chemogenetic stimulation of CeA CRF neurons (*red*, ①) triggers the release of CRF (*yellow*, ②) in the medial CeA, which may in turn stimulate GABA release (*blue*, ③) onto non-CRF neurons (*grey*) via the activation of CRF1 receptors located on intrinsic or extrinsic GABAergic presynaptic terminals [37]. **B.** Firing rates were recorded from CeA mCherry+ neurons. Top: red fluorescence and differential interference contrast images of patched CeA neuron. Bottom: representative current-clamp trace before and during CNO application (500 nM). Right: Firing rates are shown as mean ± s.e.m. (n=8 neurons). Data were analyzed by paired t-test: ***, p<0.001. **C-H.** Spontaneous inhibitory postsynaptic currents (sIPSCs) were recorded from medial CeA mCherry-neurons. **C.** Representative voltage-clamp traces before and during CNO application (500 nM) followed by subsequent R121919 application (1 µM). CNO increased sIPSC frequency (**D**), but did not affect sIPSC amplitude (**E**), rise time (**F**) or decay time (**G**). Data are shown as mean ± s.e.m. for the whole set of recorded neurons (n=34, black bars), as well as for the subset of neurons whose sIPSC frequency was altered 20% or more by CNO (n=19, red bars). Data were analyzed by paired t-test: **, p<0.01; ***, p<0.001. **H**. In a subset of CNO-sensitive cells (n=10), the CRF1 antagonist R121919 was applied following CNO. sIPSC frequencies are shown as mean ± s.e.m. and were analyzed by one-way ANOVA; **, p<0.01, Bonferroni *posthoc* test.

### Stimulation of CeA CRF neurons increases local GABAergic transmission via activation of CRF1 receptors

We first evaluated the possibility that CeA CRF neurons release CRF in the medial CeA, where both CRF and ethanol are known to increase GABA release via the activation of CRF1 receptors [37]. Voltage-clamp recordings of spontaneous inhibitory postsynaptic currents (sIPSC) were obtained from mCherry-negative neurons (n=34 neurons from 11 mice) located in the medial CeA before and during application of CNO (500 nM). Overall, CNO produced a significant increase in sIPSC frequency (Fig. 3C-D, t_33_=-3.3, p=0.002), but had no effect on amplitude (Fig. 3E, t_33_=-1.7, p=0.10), rise time (Fig. 3F, t_33_=-0.3, p=0.76) or decay time (Fig. 3G, t_33_=-0.3, p=0.79). However, the magnitude of the response to CNO was variable between cells. We categorized neurons as CNO-sensitive (n=19, red bars in Fig. 3D-G) if CNO elicited a change ≥ 20% from baseline sIPSC frequency (t_18_=-7.0, p<0.0001). In this subset of neurons, there was still no effect of CNO on sIPSC amplitude (t_18_=-1.6, p=0.12), rise time (t_18_=-0.2, p=0.86) and decay time (t_18_=-0.2, p=0.87).

In a subset of CNO-sensitive neurons (n=10), the CRF1 antagonist R121919 (1 μM) was applied after CNO to determine whether the effect of CNO on sIPSC frequency was mediated by CRF1 activation (see Fig. 3A for schematic representation of tested hypothesis). There was a significant main effect of treatment (Fig. 3H, F_2,18_=6.1, p=0.01) reflecting the increase in sIPSC frequency induced by CNO and partial reversal of the effect of CNO by R121919. *Posthoc* analysis reveal a significant difference between baseline and CNO (p=0.009), but not for other pairwise comparisons.

Altogether, these data indicate that chemogenetic stimulation of mouse CeA CRF neurons increases GABA release onto local medial CeA neurons and that this effect is at least partially mediated by CRF1 activation.

### Stimulation of CeA CRF neurons does not affect alcohol drinking

These results, combined with the fact that CRF1 signaling in the CeA drives excessive alcohol intake in rodent models of binge-like drinking and dependence [15–17], led us to hypothesize that chemogenetic stimulation of CeA CRF neurons may promote voluntary alcohol consumption. To test this hypothesis, mice were given access to ethanol (15% v:v)/water two-bottle choice (2BC) during 2-h daily sessions and the effect of CNO on ethanol intake was tested under different conditions (see Fig. 4A for experimental timeline).

**Figure 4.**
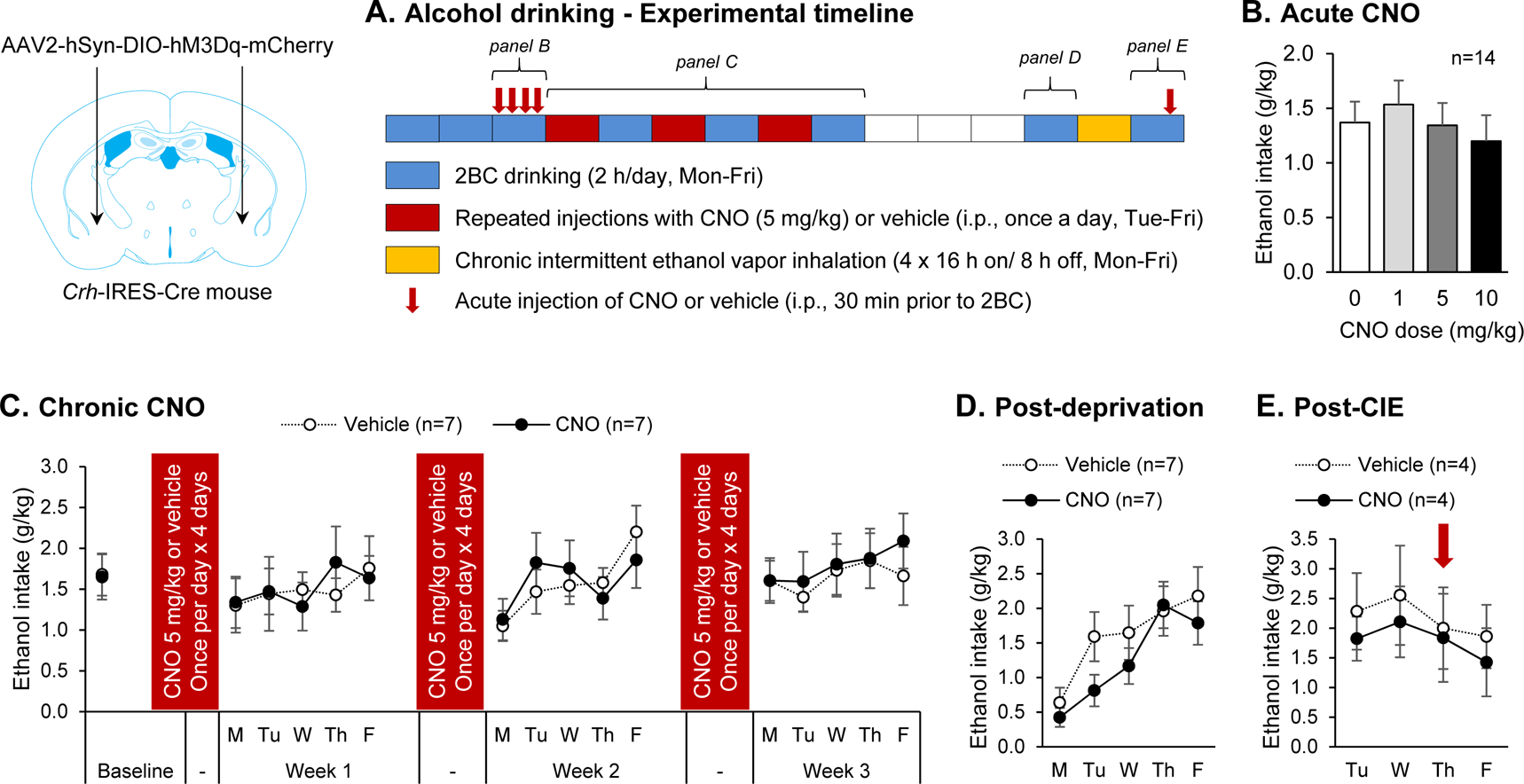
Chemogenetic stimulation of CeA CRF neurons does not affect alcohol drinking. *Crh*-IRES-Cre male mice were injected with a Cre-dependent hM3Dq-encoding vector in the anterior CeA and were tested for voluntary ethanol intake under different experimental conditions. **A.** Experimental timeline. Each box represents one week, the color code indicates the experimental procedure conducted during that week. Two-bottle choice (2BC) drinking sessions were conducted Mon-Fri, starting at the beginning of the dark phase and lasting 2 h (*blue boxes*). Mice were given ten baselining sessions prior to testing the acute effect of CNO (0, 1, 5 and 10 mg/kg, i.p., 30-min pretreatment) according to a within-subject Latin-square design over four consecutive days (*red arrows*; data shown in panel **B**). An additional 2BC session without pretreatment was conducted and mice were then split in two groups exhibiting equivalent baseline ethanol intake, which were repeatedly injected with either CNO (5 mg/kg) or vehicle. Weeks of CNO or vehicle administration (once per day, Tue-Fri; *red boxes*) were alternated with weeks of 2BC drinking sessions (Mon-Fri, as described above; *blue boxes*; data shown in panel **C**) for a total of 3 rounds. Mice were then given a 3-week ethanol deprivation period (*white boxes*), after which 2BC sessions were resumed for a week (*blue box*; data shown in panel **D**). Next, the mice were exposed to four cycles of chronic intermittent ethanol exposure (16-h ethanol vapor inhalation followed by 8-h air inhalation, Mon-Fri; *yellow box*). The mice were then returned to their home cages and 2BC sessions resumed four days later (Tue-Fri; *blue box*; data shown in panel **E**). On the third session (Thu), CNO (5 mg/kg) or vehicle was administered 30 min prior to the session (*red arrow*). **B-E.** Ethanol intake is expressed in g ethanol per kg body weight in 2-h session. Data are shown as mean ± s.e.m. Number of mice per group is shown in the legend of each graph.

Acute CNO (1, 5 and 10 mg/kg) administration (30 min prior to drinking session) did not affect ethanol intake (Fig. 4B, F_3,39_=1.6, p=0.21). We then attempted to replicate the pattern of CRF release in the CeA elicited by chronic intermittent ethanol (CIE) exposure. In the CIE-2BC model, weeks of 2BC (Mon-Fri) are alternated with weeks of CIE consisting of four 16-h periods of ethanol vapor inhalation separated by 8-h periods of air inhalation (Mon-Fri) [4]. Extracellular CRF levels are known to gradually increase in the amygdala of rats withdrawn from chronic alcohol exposure starting about 6 h after the onset of withdrawal [12]. Accordingly, to replicate the predicted pattern of CRF release experienced by CIE-exposed mice, we injected CNO (5 mg/kg) once per day (Tue-Fri) and conducted 2BC sessions on alternate weeks (Fig. 4A). In the CIE-2BC model, CIE-exposed mice typically start increasing their voluntary ethanol intake after 1-3 weeks of CIE exposure [4, 38]. However, alternating weeks of repeated CNO administration with weeks of 2BC for 3 rounds did not affect ethanol intake compared to mice repeatedly injected with vehicle (Fig. 4C). A two-way ANOVA of weekly average intakes revealed no effect of time (F_3,36_=1.4, p=0.26), no effect of treatment (F_1,36_=0.01, p=0.91) and no time x treatment interaction (F_3,36_=0.3, p=0.86). We examined whether repeated CNO administration may produce a delayed effect but there was again no effect of treatment on ethanol intake following 3 weeks of deprivation (Fig. 2D, t_12_=-1.0, p=0.33). Finally, we exposed the mice to a single week of CIE, which was not sufficient to alter ethanol intake in subsequent 2BC sessions (Tue-Fri). We then tested the ability of acute CNO (5 mg/kg) administration to sensitize the mice and increase their post-vapor ethanol intake. There was again no significant effect of CNO (Fig. 2E, t_6_=-0.2, p=0.87).

In conclusion, chemogenetic stimulation of mouse CeA CRF neurons is not sufficient to increase ethanol drinking following acute or chronic administration, or in combination with CIE exposure.

### Stimulation of CeA CRF neurons exacerbates hyponeophagia

We then examined whether chemogenetic stimulation of CeA CRF neurons may elicit anxiety-like behavior. CNO (1 or 5 mg/kg) did not affect locomotor activity in a familiar environment (Fig. 5A, F_2,22_=0.7, p=0.49). The dose of 5 mg/kg was used in all subsequent tests. In the EPM (Fig. 5B), CNO had no effect on the total distance traveled (t_12_=-0.6, p=0.56), the time spent on open arms (t_12_=1.4, p=0.19) or the number of entries on open arms (t_12_=1.3, p=0.21). In the digging assay (Fig. 5C), CNO reduced the latency to start digging (t_13_=-2.6, p=0.02) and tended to increase the total duration of digging (t_13_=1.9, p=0.08), but had no effect on marble burying (t_13_=0.2, p=0.85). In the social approach test (Fig. 5D), there was a significant main effect of compartment (F_2,24_=29.9, p<0.0001), reflecting a preference for the chamber containing a stranger mouse compared to both the center and empty cup compartments (p<0.0001 for both *posthoc* comparisons). CNO did not alter this preference, as reflected by a lack of treatment effect (F_1,12_=1.5, p=0.24) and compartment x treatment interaction (F_2,24_=0.2, p=0.80). There was also no effect of CNO on the number of transitions between compartments during habituation (vehicle: 26.1 ± 5.1; CNO: 22.7 ± 1.1; t_12_=-0.7, p=0.53) or in presence of the stranger mouse (vehicle: 33.9 ± 3.0; CNO: 36.1 ± 2.4; t_12_=0.6, p=0.56), in agreement with the lack of effect of CNO on locomotor activity.

**Figure 5.**
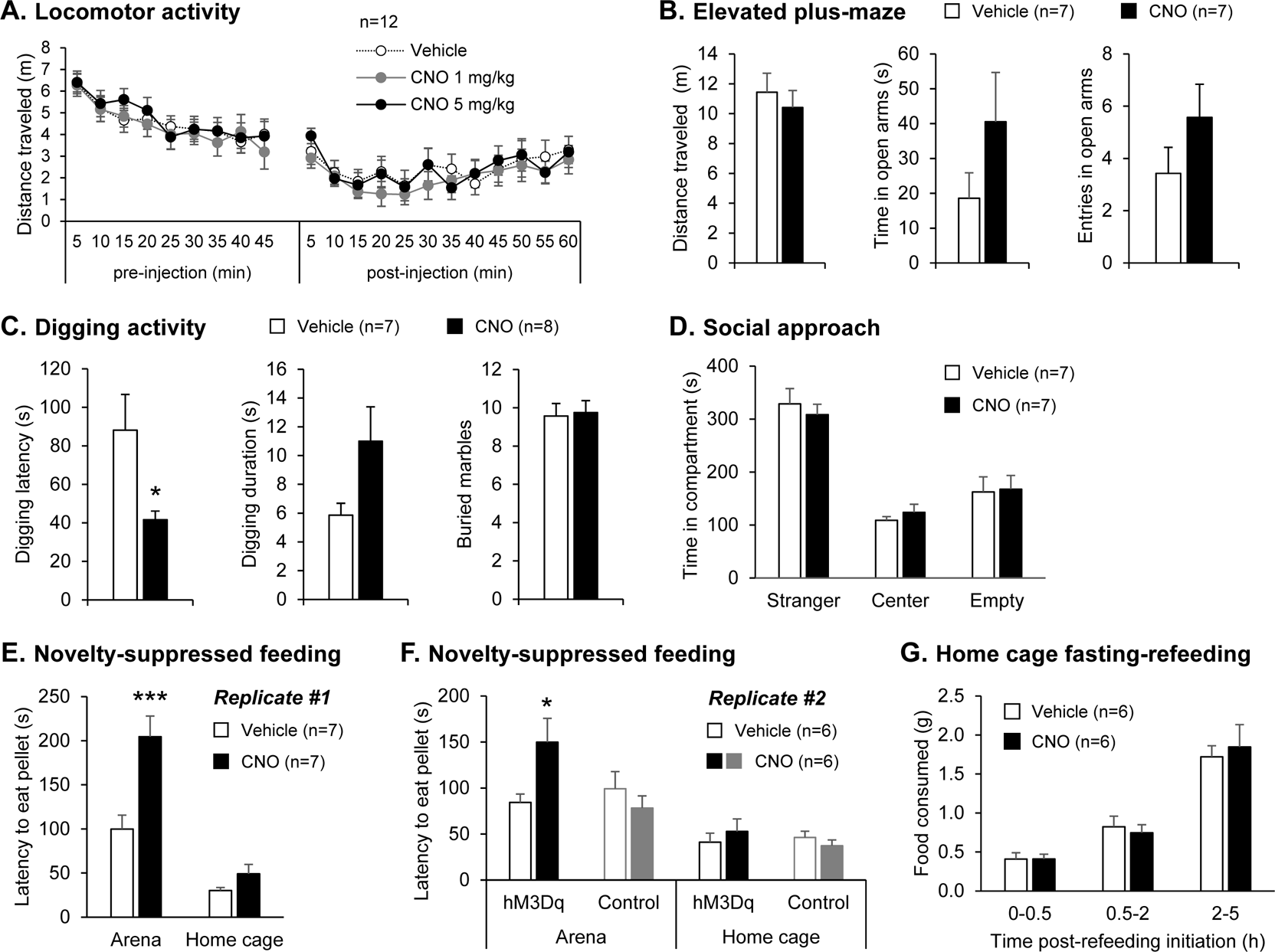
Chemogenetic stimulation of CeA CRF neurons exacerbates hyponeophagia without altering other affective responses nor appetite. *Crh*-IRES-Cre male mice were injected with a Cre-dependent hM3Dq-encoding vector in the anterior CeA. **A.** Locomotor activity 45 min prior and 60 min following i.p. injection of vehicle or CNO. **B-G.** Mice were tested in assays probing affective behavior 30 min after i.p. injection of saline or CNO (5 mg/kg). **B.** CNO did not affect open arm exploration in the elevated plus maze. **C.** CNO reduced the latency to start digging and tended to increase digging duration but did not affect marble burying. **D.** CNO did not affect the preference for social interaction. **E-F.** CNO increased the latency to start feeding in an anxiogenic arena, but not in the home cage. This effect of CNO was replicated in an independent cohort and was not observed in control mice with no hM3Dq expression. **G.** CNO did not alter the amount of food consumed following 24-h food deprivation. Data are shown as mean ± s.e.m. * indicates significant differences between Vehicle- and CNO-injected mice, ^#^ indicates significant differences between compartments (D) or environment (E). * or ^#^, p<0.05; *** or ^###^, p<0.001. Number of mice per group is shown in the legend of each graph.

In the novelty-suppressed feeding test (Fig. 5E), it took longer for the food-deprived mice to start eating in the arena than in their home cage (main effect of environment: F_1,12_=57.1, p<0.0001) and CNO selectively prolonged the arena latency (environment x treatment interaction: F_1,12_=8.3, p=0.01; *posthoc* comparison Arena-Vehicle *vs* Arena-CNO: p=0.0006; Home cage-Vehicle *vs* Home cage-CNO: p=1.00). We replicated this finding in an independent cohort, which also included control mice (Cre-dependent tdTomato expression, no hM3Dq). In this second replicate (Fig. 5F), CNO again increased the latency to start eating the pellet in the arena in hM3Dq-expressing mice (t_10_=2.4, p=0.04), but not in Control mice (t_6_=-0.9, p=0.39). To verify that the anxiogenic-like effect of CNO in the novelty-suppressed feeding test was not related to reduced appetite, we further examined the effect of CNO on feeding in a home cage setting. CNO had no effect on food consumption after 24-h deprivation at any of the time points examined (Fig.3G, F_1,10_=0.01, p=0.92).

In summary, chemogenetic stimulation of mouse CeA CRF neurons promotes hyponeophagia without altering appetite. The anxiogenic-like effect of this manipulation is not detected in other assays of anxiety-like behavior.

## Discussion

Our study shows that chemogenetic stimulation of mouse CeA CRF neurons increases GABA release onto medial CeA neurons and exacerbates hyponeophagia but does not affect voluntary alcohol consumption. These findings indicate that hyperactivity of CeA CRF neurons may contribute to elevated CeA GABA levels and anxiety-like behavior in CIE-exposed mice [8, 39–41] but is not sufficient to drive ethanol intake escalation.

There was no effect of CeA CRF neuron stimulation on the overall activity of other CeA neurons (as indexed by c-Fos expression in mCherry-negative CeA neurons). This observation contrasts with the effect of the same manipulation in *Crh*-Cre rats, where robust c-Fos induction was also detected in non-CRF neurons of the medial and lateral CeA and was mediated by CRF1 activation [28]. This discrepancy could relate to the differential anatomy of the CRF system, with the dense cluster of CRF neurons at the junction of the IPAC and anterior CeA being unique to mice and possibly exerting a stronger inhibitory control over the remainder of the CeA than the more posterior CRF neurons. Consistent with this hypothesis, the connectivity of CeA CRF neurons is species-specific, as local synapses are predominant in mice [42], while long-range projections to the BNST, lateral hypothalamus and parabrachial nucleus are robust in rats [29, 31]. Alternatively, the discrepancy may stem from a differential excitatory/inhibitory signaling balance between the two species, whereby a higher ratio of GABA to CRF may be released in mice compared to rats, inhibitory neuromodulators may be co-released to a larger extent in mice, or inhibitory signaling may override excitatory signaling via mouse-specific postsynaptic mechanisms.

The penetrance of Cre activity in CeA CRF neurons was lower in *Crh*-IRES-Cre mice (64%) than in *Crh*-Cre rats (99%), which may also explain differential experimental outcomes in the two species. However, our quantification by *in situ* hybridization may have been less sensitive than the method used in rats (immunohistochemistry, confocal imaging at 63x and 3D reconstruction) [28]. Previous characterization of the distribution of Cre activity in the amygdala of *Crh*-IRES-Cre;Ai14 mice by immunohistochemistry concluded that the pattern “largely recapitulated” the distribution of CRF, although no quantification was presented [43]. We found that the fidelity of Cre activity for CeA CRF neurons was reasonably high (86% - which may again be an underestimate), confirming that the cellular and behavioral effects of CNO reported in the present study result, for the most part, from the activation of CeA CRF neurons.

Chemogenetic stimulation of CeA CRF neurons triggered GABA release onto a subset of medial CeA neurons, as reflected by increased sIPSC frequency in response to CNO. This result is consistent with the ability of optogenetic stimulation of CeA CRF terminals to evoke IPSCs in a subset of CeA neurons in *Crh*-Cre rats and mice [28, 42]. Combined with the lack of activation of mCherry-negative CeA cell bodies we observed with c-Fos labeling, our data indicate that mouse CeA CRF neurons stimulate GABA release from their own terminals (given that CeA CRF neurons are GABAergic [44]) or possibly from extrinsic GABA afferents, but not from local non-CRF GABA interneurons as may be the case in rats (see discussion above). The ability of R121919 to partially reverse this increase in GABA release is consistent with previous work showing that exogenous CRF acts at presynaptic CRF1 receptors to increase GABA release in the mouse CeA [37].

The effect of chemogenetic stimulation of CeA CRF neurons on anxiety-like behavior appears to be species- and assay-dependent. While this manipulation did not affect mouse EPM behavior in the present study, it reduces open arm exploration in rats [29, 30]. Previous studies in *Crh*-IRES-Cre mice also showed that chemogenetic activation of CeA CRF neurons increases immobility in the EPM (although this may not necessarily translate into reduced time spent on open arms) and optogenetic activation of the CeA CRF projection to the locus coeruleus reduced exploration of open sections of an elevated zero-maze [45, 46]. We therefore anticipated that activating CeA CRF neurons would reduce open arm exploration in the EPM, but it had no effect. A floor effect may have prevented us from detecting increased anxiety-like behavior given the limited time vehicle-injected mice spent on the open arms. It is also possible that other projections of CeA CRF neurons oppose the anxiogenic-like effect of the coerulear projection, resulting in a lack of EPM phenotype in the present study.

It is important to note, however, that mice and rats can manifest negative affect in different ways, even when undergoing a similar aversive experience. For instance, while it has been repeatedly shown that rats withdrawn from chronic alcohol exposure exhibit anxiety-like behavior in the EPM [47–54], a similar phenotype has been difficult to detect in mice using the same assay. However, negative affect can be captured in alcohol-withdrawn mice using alternative assays. Most relevant to the present study, C57BL/6J males display increased digging activity and exacerbated hyponeophagia, but unaltered light-dark box exploration and social interaction, when tested 3-10 days into withdrawal from CIE [8, 41, 55–57]. The latter data are consistent with our observation that chemogenetic stimulation of CeA CRF neurons in mice exacerbates hyponeophagia and tends to increase digging activity but has no effect in the EPM and social approach tests. Altogether, this phenotypic overlap indicates that CeA CRF neurons may drive negative affect during CIE withdrawal in both rats and mice.

It is worth noting that CNO had a weaker effect than CIE withdrawal on digging activity. In contrast to CIE, it did not affect marble burying, a less sensitive measure of digging activity [8, 58]. This observation suggests that additional neural circuits contribute to increased digging activity in CIE-withdrawn mice.

We had hypothesized that CeA CRF neurons would not only drive negative affect but also increase alcohol drinking by mimicking negative reinforcement-driven escalation of alcohol self-administration elicited by CIE [59]. However, neither acute nor chronic chemogenetic stimulation of CeA CRF neurons altered alcohol intake in mice given limited access to alcohol. This observation contrasts with the ability of optogenetic inhibition of the same neuronal population to selectively reduce escalated alcohol self-administration in CIE-exposed rats [31]. This discrepancy suggests that CeA CRF neurons are *necessary* but not *sufficient* to drive alcohol intake escalation in dependent animals, i.e. that additional neural circuits are required for negative affect to influence the motivation to consume alcohol.

Aside from fear learning and anxiety, CeA neurons have been implicated in feeding control. Specifically, lateral CeA PKCδ-positive neurons suppress food intake [60]. In the novelty-suppressed feeding test, we found that chemogenetic stimulation of CeA CRF neurons increased the latency of food-deprived mice to start eating in an anxiogenic arena but did not affect their home cage feeding. This observation is consistent with the minimal overlap between mouse CeA populations expressing PKCδ versus CRF [42, 61]. Interestingly, CRF suppresses feeding when administered in the brain and CRF signaling mediates the anorexigenic effect of several stressors (see [62] for review). While CeA CRF neurons can be recruited by stress ([63, 64], but see [65]), our findings indicate that their activity is not sufficient to inhibit feeding.

Altogether, our results indicate that excitation of CeA CRF neurons, which is known to happen during alcohol withdrawal [31, 66], is sufficient to produce negative affect but not to increase the motivation to consume alcohol. These data suggest that alcohol drinking escalation in animals made dependent to alcohol is not solely driven by negative reinforcement associated with activation of CeA CRF neurons. Future work will be needed to identify other mechanisms contributing to alcohol drinking escalation in dependence, including additional neural circuits releasing CRF in the CeA as well as non-CRF signaling pathways that may potentiate the influence of CRF transmission on alcohol drinking.

## Acknowledgments

We wish to thank Sophia Zhu and Tanvi Shah for their assistance with immunohistochemistry. This work was supported by the following grants from the National Institutes of Health: AA024198 (CC), AA026685 (CC), AA006420 (CC and MR), AA021491 (MR), AA015566 (MR), AA023002 (MAH), and AA024952 (HS).

