## Supplementary Methods for "Chemogenetic stimulation of mouse central amygdala corticotropin-releasing factor neurons: Effects on cellular and behavioral correlates of alcohol dependence"

***Animals***

C57BL/6J males were purchased from the Jackson Laboratories (stock #000664) at 8 weeks of age. *Crh*-IRES-Cre male breeders were obtained from The Jackson Laboratory (B6(Cg)Crh^tm1(cre)Zjh^/J, stock # 012704, [1]) and were mated with C57BL/6J females from The Scripps Research Institute rodent breeding colony to generate the heterozygous mice used for experimentation. Backcross breeders were introduced every 1-2 years to prevent genetic drift. Two male offspring from *Crh*-IRES-Cre and Ai9 (B6.Cg-Gt(ROSA)26Sor^tm9(CAG-tdTomato)Hze^/J, The Jackson Laboratory, stock #007909, [2]) mating were kindly provided by the laboratory of Dr. Lisa Stowers (The Scripps Research Institute, La Jolla, CA) and used for double fluorescent *in situ* hybridization.
Mice were maintained on a 12 h/12 h light/dark cycle. Food (Teklad LM-485, Envigo) and acidified or reverse osmosis purified water were available *ad libitum* except for a 24-h period of food deprivation before the novelty-suppressed feeding and home cage fasting-refeeding tests. Sani-Chips (Envigo) were used for bedding substrate. Mice were at least 10 weeks old at the time of surgery. Only heterozygous males were used for experiments. They were group-housed except for alcohol drinking experiments. All procedures adhered to the National Institutes of Health Guide for the Care and Use of Laboratory Animals and were approved by the Institutional Animal Care and Use Committee of The Scripps Research Institute.

***Drugs***

Clozapine-N-oxide (CNO) was obtained from Sigma (C0832) for electrophysiological recordings and Enzo Life Sciences Inc. (BML-NS105-0025) for behavioral assays and c-Fos induction. CNO was dissolved in dimethyl sulfoxide (DMSO) and diluted in 0.9% saline for intraperitoneal (i.p.) injection (10 mL/kg body weight). Vehicle solution contained 0.5% DMSO and CNO was injected at a dose of 1 or 5 mg/kg. Chloral hydrate was purchased from Sigma-Aldrich, dissolved in sterile water at a concentration of 35% (w:v) and injected i.p. (10 mL/kg body weight). DNQX (6,7-dinitroquinoxaline-2,3-dione, 10 µM), AP-5 (DL-2-amino-5-phosphonovalerate, 50 µM) and CGP 55845A (1 µM) were purchased from Tocris Bioscience. R121919 (1µM) was supplied by Neurocrine Biosciences, Inc.

***Viral vectors***

Adeno-associated viral serotype 2 (AAV2) vectors encoding the hM3Dq excitatory designer receptor [3] fused to the red fluorescent protein mCherry, under the control of the human synapsin promoter and in a Cre-dependent manner (AAV2-hSyn-DIO-hM3Dq-mCherry [4]), were obtained from the Vector Core at the University of North Carolina (UNC) at Chapel Hill (lot AV4499G, titer 5.1 x 10^12^ vg/mL; lot AV4499H, titer 2 x 10^12^ vg/mL) or from Addgene (lot v6233, titer 4.6 x 10^12^ vg/mL). An AAV2 vector encoding cytoplasmic tdTomato and presynaptic (synaptophysin-fused) EGFP [5, 6] downstream of the human synapsin promoter and upon Cre recombination (AAV2-hSyn-FLEX-tdTomato-T2A-SypGFP [7]) was obtained from UNC Vector Core (lot AV7138, titer 2.7 x 10^12^ vg/mL) and used as negative control for hM3Dq expression.

***Experimental cohorts***

*In situ* hybridization data were collected from a cohort of C57BL/6J mice subjected to chronic alcohol drinking (n=15). For chemogenetic experiments, a first cohort of *Crh*-IRES-Cre mice was used for electrophysiological recordings (n=12, injected with viral stock AV4499G). A second cohort was tested in the elevated plus-maze (EPM), social approach, and novelty-suppressed feeding assays, with at least one week between tests; these mice were then single-housed and tested for alcohol drinking (n=7, AV4499G; n=7, v6233). A third cohort was tested for digging and marble burying, and their brains were used to quantify c-Fos induction one week later (n=15, v6233). A fourth cohort was used to confirm the phenotype observed in the novelty-suppressed feeding test (n=12, AV4499H) and include negative control mice (n=8, AV7138); these mice were also tested for locomotor activity and fasting-refeeding in the home cage.

***In situ hybridization***

Mice were quickly decapitated and brains were snap-frozen in isopentane. Ten series of 20-μm coronal sections were sliced in a cryostat and directly mounted on Superfrost slides and stored at -80°C. Chromogenic *in situ* hybridization (CISH) was conducted as described in [8]. A pBlueScript plasmid containing the rat *Crh* cDNA (1.1 kb) was donated by Dr. Kelly Mayo (Northwestern University, Evanston, IL). Digoxigenin (DIG)-labeled riboprobes were synthesized using a kit (Roche, Indianapolis, IN). Sections were post-fixed in PFA 4%, and then acetylated in 0.1 M triethanolamine pH 8.0, acetic acid 0.2%. Following washes in salt sodium citrate (SSC) 2x, sections were dehydrated and defatted in a graded ethanol/chloroform series. Pre-hybridization and hybridization were performed at 70°C in a buffer containing 50% formamide, SSC 2x, Ficoll 0.1%, polyvinylpyrrolidone 0.1%, bovine serum albumin 0.1%, sheared salmon sperm DNA (0.5 mg/mL) and yeast RNA (0.25 mg/mL). Probes were diluted in the hybridization buffer (800 ng/mL) and incubated overnight on slides. Post-hybridization washes were performed in 50% formamide, SSC 2x and Tween-20 0.1%. Sections were then blocked for 1 h and incubated with anti-DIG antibody overnight at 4°C (Roche, 1:2000) in MABT buffer (0.1 M maleic acid pH 7.5, 0.15 M NaCl, Tween-20 0.1%) containing 10% normal goat serum. Following washes in MABT and incubation in detection buffer (0.1 M Tris-HCl pH 9.5, 0.1 M NaCl, 0.05 M MgCl2, Tween-20 0.1%), the reaction with NBT-BCIP was allowed to develop in the dark for 24 h at room temperature. Slides were rinsed, air dried and mounted in DPX (Sigma, St-Louis, MO). Sections containing the CeA at three anteroposterior levels (bregma -0.8 mm, -1.2 mm and -1.6 mm) were imaged at 5x (one image per hemisphere) using a Zeiss Axiophot microscope equipped with a QImaging Retiga 2000R color digital camera and QCapture software. Images were converted to grayscale and optical density of the CISH signal in the CeA was analyzed using NIH Image J software.
Double fluorescent *in situ* hybridization was conducted similarly, with the following modifications. A 685-bp fragment of *tdTomato* cDNA (monomer sequence) was subcloned in pBlueScript II and linearized for synthesis of a fluorescein-labeled probe. Sections were pretreated with H_2_O_2_ 3% prior to acetylation. After hybridization, the *Crh* and *tdTomato* probes were detected consecutively by immunostaining with horseradish peroxidase-conjugated antibodies (Roche, 1:200) incubated in TNT buffer (0.1 M Tris-HCl pH 7.5, 0.15 M NaCl, Tween-20 0.1%) containing 1x blocking reagent (Roche), followed by tyramide signal amplification (Perkin-Elmer, 1:50, TSA-Cy3 and TSA-Fluorescein, respectively). Images were captured using a Keyence BZ-X700 fluorescence microscope. Red, green, and co-labeled cell bodies were counted in Image J.

***Stereotaxic surgery***

Mice were anesthetized with isoflurane, placed in a stereotaxic frame (David Kopf Instruments, model 940), and injected bilaterally into the anterior part of the CeA (AP -0.9 mm from bregma, ML ± 3.0 from the midline, DV -4.5 mm from the skull) with 1 μL of vector using a 22G double guide cannula and 28G single injectors projecting 5 mm (Plastics One). The vector was infused at a rate of 0.1 μL/min for 10 min using a dual syringe pump (Harvard Apparatus). The injectors were left in place for an additional 10 min and slowly retracted to minimize backflow. The scalp was sutured using surgical thread. Mice were left undisturbed for at least three weeks post-injection prior to recording or behavioral testing.

***Electrophysiological recordings***

Mice were exposed to brief anesthesia (3-5% isoflurane) after which brains were rapidly extracted, placed in ice-cold sucrose solution containing (in mM): sucrose 206.0; KCl 2.5; CaCl_2_ 0.5; MgCl_2_ 7.0; NaH_2_PO_4_ 1.2; NaHCO_3_ 26; glucose 5.0; HEPES 5, and coronally sectioned (300 μm) on a Leica VT1000S (Leica Microsystems). After sectioning, slices were incubated in an oxygenated (95% O_2_/5% CO_2_) artificial cerebrospinal fluid (aCSF) solution containing (in mM): NaCl 130, KCl 3.5, NaH2PO4 1.25, MgSO4 1.5, CaCl2 2, NaHCO3 24, glucose 10 for 30 min at 37°C, followed by 30 min equilibration at room temperature (21-22°C). Recordings were made with pipettes (3-6 MΏ; King Precision Glass) filled with an internal solution containing (in mM): KCl 145; EGTA 5; MgCl_2_ 5; HEPES 10; Na-ATP 2; Na-GTP 0.2 or an internal solution containing (in mM) Kgluconate 145; EGTA 5; MgCl_2_ 5; HEPES 10; Na-ATP 2; Na-GTP 0.2, coupled to a Multiclamp 700B amplifier (Molecular Devices), acquired at 10 kHz, low-pass filtered at 2-5 kHz, digitized at 20 kHz (Digidata 1440A; Molecular Devices), acquired and stored using pClamp 10 software (Axon Instruments). Series resistance was continuously monitored 10 mV pulses; neurons with series resistance >15 MΩ or >20% change in resistance during recording were excluded from final analysis. CeA neurons containing mCherry were identified and differentiated from unlabeled neurons using fluorescent optics and brief (<2 s) episcopic illumination. Electrophysiological properties of cells were determined by pClamp 10 Clampex software online during voltage-clamp recording using a 10-mV pulse delivered after breaking into the cell. Drugs were applied either by bath or Y-tube application for local perfusion. Recordings (V_hold_= -60 mV) were performed in the presence of the glutamate receptor blockers 6,7-dinitroquinoxaline-2,3-dione (DNQX, 20 µM) and DL-2-amino-5-phosphonovalerate (DL-AP5, 30 µM) and the GABA_B_ receptor antagonist CGP55845A (1 µM).

***Immunohistochemistry***

Mice were anesthetized with chloral hydrate and perfused with cold phosphate buffered saline (PBS) followed by 4% paraformaldehyde (PFA). Brains were dissected and immersion fixed in PFA for 24 h at 4°C, cryoprotected in sterile 30% sucrose in PBS at 4°C or until brains sank and stored at ‑80°C. Coronal 35-µm thick brain sections were sliced with a cryostat, collected in 5 series in PBS containing 0.01% sodium azide, and stored at 4°C. Throughout the immunostaining procedure, sections were gently agitated and steps were performed at room temperature unless noted otherwise. Sections were first washed in PBS for 10 min, then blocked for 1 h in PBS containing 0.3% triton X-100, 1mg/ml bovine serum albumin (BSA) and 5% normal goat serum (NGS). Primary antibodies (rabbit anti-mCherry, Abcam ab167453, 1:5,000; guinea-pig anti-c-Fos, Synaptic Systems 226004, 1:2,000) were diluted in PBS containing 0.5% Tween-20 and 5% NGS and incubated at 4°C overnight. Next, sections were washed in PBS (10 min, 3 times) followed by a 1-h incubation with secondary antibodies (AlexaFluor 568 goat anti-rabbit, Thermo Fisher Scientific A-11011, 1:500; AlexaFluor 488 goat anti-guinea pig, Thermo Fisher Scientific A-11073, 1:500) diluted in PBS. Sections were then washed (10 min, 3 times) and air-dried, and coverslips were mounted using DAPI-containing Vectashield Hardset medium (Vector Laboratories, H1500). Images were captured using a Zeiss LSM 710 confocal laser scanning microscope or a Keyence BZ-X700 fluorescence microscope. mCherry labeling was used to verify the location of hM3Dq-expressing cell bodies in mice subjected to behavioral testing. c-Fos labeling was used to confirm the excitatory effect of CNO in hM3Dq-expressing cells. In the latter experiment, mice were injected with vehicle or CNO (5 mg/kg, i.p.) 90 min prior to perfusion, and counts of single- and double-labeled cells were obtained from both hemispheres (number of images analyzed in each mouse: vehicle (n=7), 6.7 ± 0.5 images; CNO (n=6), 6.5 ± 0.8 images).

***Behavioral testing***

All behavioral testing was conducted during the dark phase of the light/dark cycle.

***Alcohol drinking***

Mice were single-housed and food pellets were placed in the bedding instead of the food hopper throughout the duration of the experiment. Two-bottle choice (2BC) drinking sessions were conducted Mon-Fri, starting at the beginning of the dark phase and lasting 2 h. During these sessions, the home cage water bottle was replaced with two 50-mL conical tubes fitted with a rubber stopper and sipper tube assembly and filled with acidified water or ethanol 15% v:v, respectively. The positions of the water and ethanol bottles were alternated every day and bottles were weighed at the end of each session. Body weights were measured on a weekly basis to calculate ethanol intake (g/kg).

For *Crh* expression data, C57Bl/6J mice were subjected to nine weeks of 2BC and brains were collected four weeks after their last drinking session.

For chemogenetic manipulation, the experimental timeline is explained in Figure 4A. Mice were given ten baselining sessions prior to testing the effect of CNO (0, 1, 5 and 10 mg/kg, i.p., 30-min pretreatment) according to a within-subject Latin-square design over four consecutive days. We then tested the delayed effect of repeated CNO administration on subsequent ethanol intake in an attempt to mimic the effect of repeated alcohol withdrawal in mice subjected to chronic intermittent ethanol inhalation (CIE, [9]). An additional 2BC session without pretreatment was conducted and mice were then split in two groups exhibiting equivalent baseline ethanol intake, which were repeatedly injected with either CNO (5 mg/kg) or vehicle. Weeks of CNO or vehicle administration (once per day, Tue-Fri) were alternated with weeks of 2BC drinking sessions (Mon-Fri, as described above) for a total of 3 rounds. Mice were then given a 3-week ethanol deprivation period, after which 2BC sessions were resumed for a week. Next, the mice were exposed to four cycles of CIE, as previously described [10, 11]. Briefly the mice were subjected to four 16-h periods of ethanol vapor inhalation separated by 8-h periods of air inhalation (Mon-Fri). Six mice (vehicle, n=3; CNO, n=3) were excluded from the experiment at this point due to fighting-induced wounds. Each intoxication period was initiated with an intraperitoneal injection of ethanol (1.5 g/kg) and pyrazole (68 mg/kg). Average serum ethanol concentration during intoxication was 195.8 ± 15.9 mg/dL, as measured by gas chromatography and flame ionization detection (Agilent 7820A). The mice were then returned to their home cages and 2BC sessions resumed four days later (Tue-Fri). On the third session (Thu), CNO (5 mg/kg) or vehicle was administered 30 min prior to the session.

***Locomotor activity.*** Mice were placed in individual cages lined with bedding. The same cage was used across days for each individual mouse. Mice were first habituated to the setup for 2 h on two consecutive days. On testing days, locomotor activity was recorded for 45 min, injections were administered, and videotracking was then resumed for another 60 min. CNO (0, 1 and 5 mg/kg) was administered according to a within-subject Latin-square design over three consecutive days. The horizontal motion of each mouse was videotracked using ANY-maze software (Stoelting) and the distance traveled during 5-min bins was analyzed.

***Elevated plus maze (EPM)***

Testing was conducted under dim lighting (20 lux on open arms). The apparatus was made of black acrylic and the dimensions were as follows: 30-cm long x 5-cm wide arms, 15-cm high walls for the closed arms, 30-cm elevation. The mice were habituated to the testing room for at least 1 h and injections were administered 30 min prior to the test. Each mouse was tested only once and the effect of CNO was evaluated using a between-subject design. The path of the mouse was videotracked using ANY-maze. Total distance traveled, number of entries onto open and closed arms and time spent in each area of the maze were analyzed.

***Digging and marble burying***

Testing was conducted under dim lighting (20 lux). The mouse was placed in a new, clean cage with a bedding thickness of 5 cm and no lid, and allowed to freely dig for 3 min. The number of digging bouts and total digging duration were recorded. The mouse was then removed from the cage, the bedding flattened, and 12 marbles arranged in a 4 x 3 array on top of the bedding. The mouse was reintroduced into the cage and allowed to bury the marbles for 30 min with a lid covering the cage. The number of marbles that were buried (two-thirds or more) was counted at the end of the test.

***Social approach test***

Testing was conducted under red lights. The apparatus was a rectangular box constructed from clear acrylic and containing three adjacent chambers 19 cm x 45 cm each, with 30-cm high walls. The three chambers were separated by dividing walls made from clear acrylic with openings between the central chamber and each side chamber. Mice were first habituated to the apparatus during 5 min, with free access to all three compartments of the apparatus. An unfamiliar mouse (C57BL/6J male) was then placed under a small wire cup (Galaxy cup, Spectrum Diversified) in one of the side chambers and an empty cup was placed in the other side chamber. The test mouse was then allowed to explore all three chambers for 10 min. The time spent in each chamber (all four paws) and the time the test mouse spent interacting with the stranger mouse were recorded manually. Placement of the stranger mouse in either of the two side chambers was randomly alternated between test mice. The number of transitions between compartments were counted during both phases.

***Novelty-suppressed feeding***

Hyponeophagia was measured by presenting regular chow to food-deprived mice. Mice were weighed and transferred to a new, clean home cage without food approximately 24 h before testing. Right before testing, mice were transferred to new, clean holding cages to clear home cages. Testing consisted of two consecutive phases: feeding in arena and feeding in home cage. The experimental arena consisted of a brightly lit (400 lux) Taconic Transit Cage (56-cm long x 40-cm wide x 18-cm deep) whose bottom was lined with 2 cm of fresh bedding. One food pellet was secured onto a platform located in the center of the arena, as described in [12]. The test mouse was placed in a corner of the arena and the latency to eat the food pellet was recorded, with a cutoff time of 10 min. The mouse was removed from the arena as soon as it started eating the food pellet and was immediately transferred to its home cage with a single food pellet of known weight (no lid on the cage, same lighting condition as the arena). The latency to eat this pellet was also recorded.

***Fasting-refeeding test***

Mice were transferred to a new, clean home cage without food. Twenty-four hours later, a pre-weighed food pellet was introduced in the cage. The water bottle and lid were put back in place. The pellet was weighed again 30 min, 2 h and 5 h later to measure amount of food consumed.
